# A novel non-invasive biomarker based on oral microbiome dysbiosis for detection of Community-Acquired Pneumonia

**DOI:** 10.1101/2022.10.25.513807

**Authors:** Ni Sun, Xuhan Zhang, Yating Hou, Ting Zhong

**Affiliations:** School of Stomatology, Jinan University, Guangzhou 510632, China

**Keywords:** Oral microbiome, Biomarker, Community-acquired pneumonia, 16S rRNA

## Abstract

**Background:** Early diagnosis of pathogenic bacteria is crucial for the treatment of community-acquired pneumonia (CAP), but conventional diagnostics are limited by sampling difficulties. Oral microbiota has also been explored as a noninvasive biomarker of lung diseases, but it’s role in CAP has been neglected. We aimed to investigate whether the oral bacteria can be novel non-invasive biomarkers for CAP.

**Methods:** Oral swab samples were collected from 29 patients with CAP and 26 healthy volunteers and characterized based on clinical parameters and 16S rRNA profiling of oral bacteria. A predict functional profiling was performed for the functional and metabolic changes in oral microbial communities.

**Results:** Oral microbial of patients with CAP had a lower diversity than healthy group. And the dominant bacteria were *Streptococcus, Prevotella* and *Neisseria* in CAP. Higher abundance of *Prevotella* (particularly *Prevotella_melaninogenica*), *Veillonella* and *Campylobacter*, and lower abundance of *Neisseria* and *Fusobacterium* were detected in CAP group. Analysis of the functional potential of oral microbiota demonstrated that the pathway involving infectious disease was overrepresented in the CAP groups relative to that in the healthy controls.

**Conclusions:** Oral microbial dysbiosis was found in patients with CAP, supporting the use of this non-invasive specimen for biomarkers of CAP.

**Highlights:**

- Oral microbial diversity was significantly lower in community-acquired pneumonia (CAP) patients than healthy controls.
- Genera *Neisseria* and *Fusobacterium* were decreased, while genera *Prevotella, Veillonella* and *Campylobacter* were increased in CAP versus healthy controls.
- Oral microbiota-based biomarkers can serve as a promising non-invasive tool for the detection of CAP.

## 1. Introduction

Community-acquired pneumonia (CAP) is the leading cause of death among infectious diseases and the third leading cause of death worldwide (1-3). As we know, the most common pathogens of CAP are *Streptococcus pneumoniae, Haemophilus influenzae* and *Staphylococcus aureus*, or viruses such as influenza viruses (4). Up to now, the cornerstone of treatment and management for patients with pneumonia is empirical antibiotic therapy (5). However, with the extensive use of antibiotics, microbiome dysbiosis and antibiotic resistance has brought a huge clinical and economic burden to CAP (6).

Conventional diagnostics of pathogen and resistance determination still rely on culture-based methods. These techniques have certain limitations, leaving correct initial antimicrobial therapy to chance (7, 8). The main microbial assessment techniques are sputum culture, blood culture, invasive technique, and urine antigen detection (5). However, these microbial assessment methods may be limited by sampling difficulties, delayed results, and difficulties in interpretation (7, 9, 10). There is a need for improved microbiological diagnostic techniques for CAP to optimize future treatment choices.

Up to now, accumulating evidence shows that oral cavity is the site of the primary source community of lung microbiota (acquired via micro aspiration and inhalation) (11, 12). Kentaro et al. observed associations between variations of oral microbiota in patients with CAP and aspiration risks (13). Jankl et al. identified the role of poor oral health and oral microbiome dysbiosis in the development of pneumonia and exacerbation of pneumonia-related complications (9). Recently, the oral microbiota has also been explored as a biomarker of lung disease. The variation in oropharyngeal microbiome of COVID-19 patients may be used as a noninvasive biomarker for dysbiosis of the pulmonary microbiome or for invasion of potential pathogens in the lungs (14).

The mouths of inpatients with CAP were more frequently colonized by respiratory pathogens (12). Oral microbial diversity may be used to develop microbiome-based biomarkers, which may have potential implications for the prevention, early diagnosis, and treatment of diseases such as pneumonia (15). But oral factors have been relatively neglected in the etiology and diagnosis of CAP (12). Little is known about the characteristics of the oral microbiota in patients with CAP and whether it can be used as a marker for early diagnosis.

Here, we used 16S rRNA gene sequencing to explore the typical profile oral bacteria in patients with CAP and found the noninvasive biomarkers for CAP.

## 2. Methods

### 2.1 Study design and sample collection

Twenty-nine consecutive CAP patients in the First Affiliated Hospital, Jinan University were enrolled in this study. CAP was defined according to the Infectious Diseases Society of America (IDSA) / American Thoracic Society (ATS) guidelines for diagnosing CAP in adults (16). This study was approved by the Human and Animal Ethics Review Committee of Jinan University, China (JNUKY-2022-002). Written informed consent was obtained from either the patients or their guardians. Finally, 55 eligible cases including 29 patients with CAP and 26 matched healthy family members were included in this study according to the recruitment process. Oral samples were collected from participants by rubbing the insides of both cheeks, palate, tongue, and teeth with a swab as described previously (9, 17). The swab containing the specimen was placed in a sterile tube and then stored at -80°C until further analysis. The blank control sample was obtained by opening the sterile tube and placing it in the sample collection environment for 5 minutes. The following basic information was collected: age, sex, BMI, underlying diseases, clinical manifestations, and oral health status.

Inclusion criteria of patients:

1. Patients with clinical and imaging diagnosis of CAP.
2. Age 18-80, with independent reading and writing skills.
3. More than 20 natural teeth.
4. Sign informed consent and voluntarily join the experiment.

Exclusion criteria of patients:

1. Suffering from serious cardiovascular diseases, tumors and other diseases that cannot be controlled by drugs, etc.
2. Patients with difficulty in completing questionnaire or oral examination.

### 2.2 DNA extraction for microbiome analysis

Genomic DNA from oral swab samples was extracted using CTAB (18). DNA integrity and size were verified by 1.0% agarose gel electrophoresis, and DNA concentrations were determined using the spectrophotometer.

### 2.3 High throughput 16S ribosomal RNA gene sequencing

16S ribosomal RNA (rRNA) gene amplification was performed using the primers(515F: GTGCCAGCMGCCGCGGTAA; 806R: GGACTACHVGGGTWTCTAAT) directionally targeting the V3 and V4 hypervariable regions of the 16S rRNA gene. To differentiate each sample and yield accurate phylogenetic and taxonomic information, the gene products were attached with forward and reverse error-correcting barcodes.

The amplicons were quantified after purification. Then, the normalized equimolar concentrations of each amplicon were pooled and sequenced on the NovaSeq6000 sequencing instrument.

### 2.4 Sequencing data analysis

QIIME 2 software was used to conduct all diversity analyses of the sequenced microbial communities. Its key plug-in is DATA 2 (19). Different from the traditional OTU-based analysis methods, DADA2 method is mainly used for noise reduction. It does not cluster with similarity but only with dereplication, which is equivalent to clustering with 100% similarity, resulting in ASVs (Amplicon sequence variants).

The Shannon index was used to assess alpha diversity; whereas weighted and unweighted UniFrac distance matrices were used to assess beta diversity, which was visualized through a principal coordinate analysis (PCoA). Beta diversity dissimilarity between groups was tested via an analysis of similarity (ANOSIM) with the R (v 3.4.1) vegan package. Bacterial taxa that differed significantly in abundance of oral microbiome between the CAP group and control group (the core shared oral bacteria) were identified via a linear discriminant analysis (LDA) effect size (LEfSe) approach (20). Significant enrichment was defined as an LDA > 3 and a p-value < 0.05.

### 2.5 Predictive functional profiling

To study the functional and metabolic changes in oral microbial communities, all ASVs were aligned into the Phylogenetic Investigation of Communities by Reconstruction of Unobserved States (PICRUSt) built-in reference database (21).

### 2.6 Statistical analysis

The SPSS software package (version 21) was used. Associations between the clinical characteristics were performed by Pearson’s Chi-square test or Fisher’s exact test. P < 0.05 was considered significant.

## 3. Results

### 3.1 Participant information

In total, 55 eligible cases including 26 healthy family numbers, 29 patients with CAP were included in this study according to the recruitment process. The basic conditions of the two groups were in good agreement. There were no significant differences in the basic information, oral hygiene habits and oral health status between the two groups. Detailed clinical data for the studied individuals were shown in Table 1.

**Table 1.**
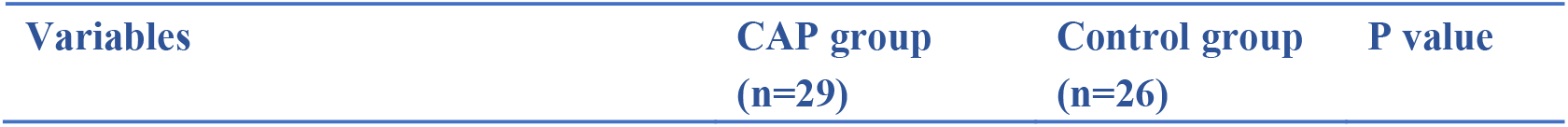
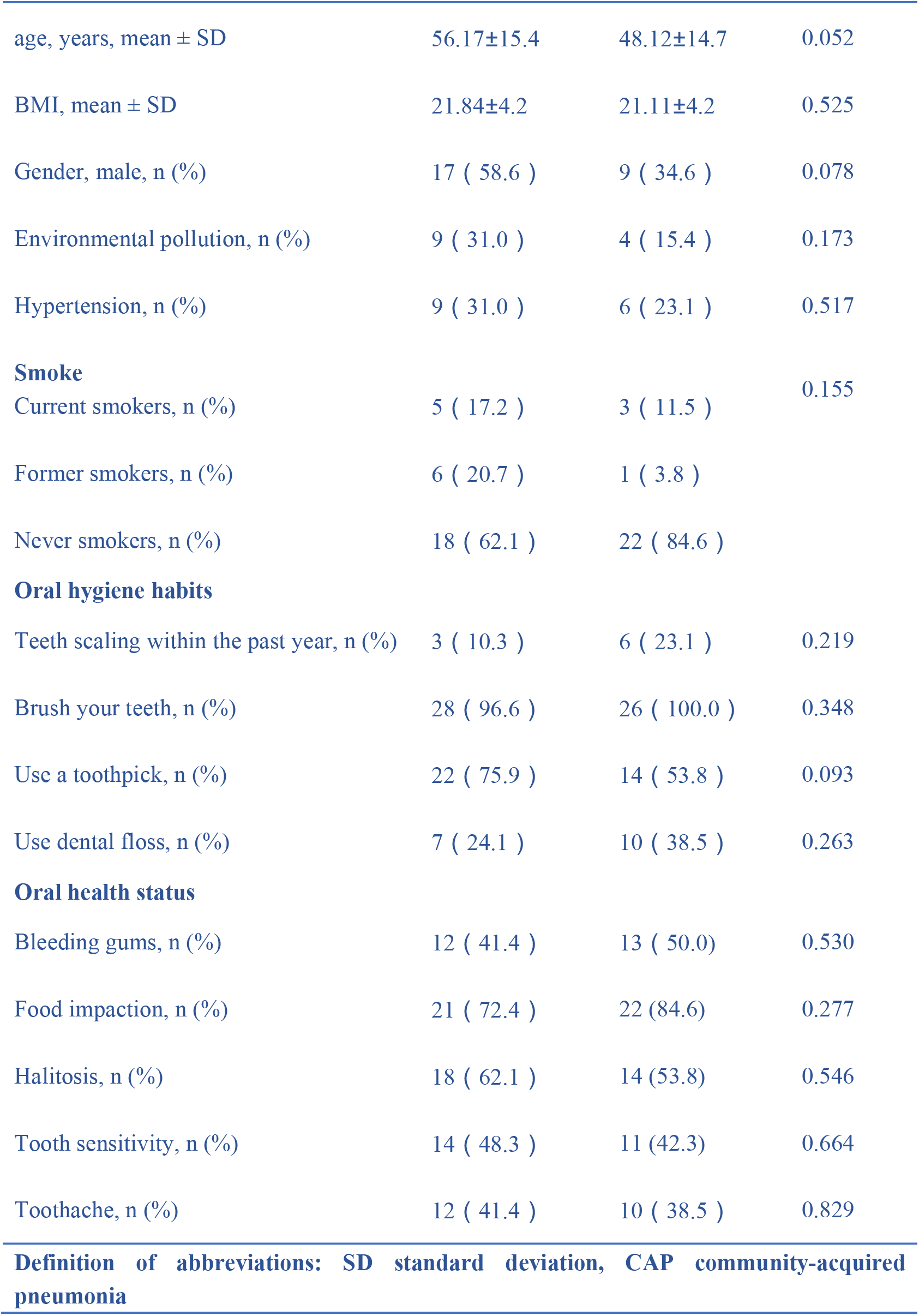
Clinical characteristics of the enrolled participants

| Variables | CAP group<br>(n=29) | Control group<br>(n=26) | P value |
| --- | --- | --- | --- |
| age, years, mean ± SD | 56.17±15.4 | 48.12±14.7 | 0.052 |
| BMI, mean ± SD | 21.84±4.2 | 21.11±4.2 | 0.525 |
| Gender, male, n (%) | 17 ( 58.6 ) | 9 ( 34.6 ) | 0.078 |
| Environmental pollution, n (%) | 9 ( 31.0 ) | 4 ( 15.4 ) | 0.173 |
| Hypertension, n (%) | 9 ( 31.0 ) | 6 ( 23.1 ) | 0.517 |
| **Smoke** |  |  | 0.155 |
| Current smokers, n (%) | 5 ( 17.2 ) | 3 ( 11.5 ) |  |
| Former smokers, n (%) | 6 ( 20.7 ) | 1 ( 3.8 ) |  |
| Never smokers, n (%) | 18 ( 62.1 ) | 22 ( 84.6 ) |  |
| **Oral hygiene habits** |  |  |  |
| Teeth scaling within the past year, n (%) | 3 ( 10.3 ) | 6 ( 23.1 ) | 0.219 |
| Brush your teeth, n (%) | 28 ( 96.6 ) | 26 ( 100.0 ) | 0.348 |
| Use a toothpick, n (%) | 22 ( 75.9 ) | 14 ( 53.8 ) | 0.093 |
| Use dental floss, n (%) | 7 ( 24.1 ) | 10 ( 38.5 ) | 0.263 |
| **Oral health status** |  |  |  |
| Bleeding gums, n (%) | 12 ( 41.4 ) | 13 ( 50.0) | 0.530 |
| Food impaction, n (%) | 21 ( 72.4 ) | 22 (84.6) | 0.277 |
| Halitosis, n (%) | 18 ( 62.1 ) | 14 (53.8) | 0.546 |
| Tooth sensitivity, n (%) | 14 ( 48.3 ) | 11 (42.3) | 0.664 |
| Toothache, n (%) | 12 ( 41.4 ) | 10 ( 38.5 ) | 0.829 |
| **Definition of abbreviations: SD standard deviation, CAP community-acquired pneumonia** |  |  |  |

### 3.2 Bacterial diversity of oral microbiota was significantly lower in patients with CAP

The oral microbiota was assessed using 16S rRNA sequencing. A total of 3,762,330 high-quality 16S rRNA gene sequences were identified, with a median read count of 68,141 (ranging from 60,123 to 76,354) per sample. After the taxonomic assignment, 4796 ASVs were obtained (<u>Table S1</u>). The species accumulation curve of all samples basically approached the asymptote, supporting the adequacy of our sampling efforts (Figure 1A). Likewise, pielou’s evenness was evaluated by rarefaction curve, exhibiting similar patterns in all samples (Figure 1B), which reflected the evenness of the distribution of species in the microbial community. Alpha diversity indexes were calculated to assess the differences in bacterial diversity between the two groups (<u>Table S2</u>). The results showed that oral microbial alpha diversity was significantly lower in the CAP group than in the healthy controls (Figure 1C-D). Moreover, the Venn diagram showed that 1832 of the total 4796 ASVs were shared between the two groups. Notably, 1080 ASVs were unique for the CAP group (Figure 1E). To display the microbiome space between samples, beta diversity was calculated using the weighted UniFrac method and the principal coordinate analysis (PCoA) was performed. The results showed a gradually separated distribution of the oral microbial communities between these two groups (Figure 1F).

**Figure 1.**
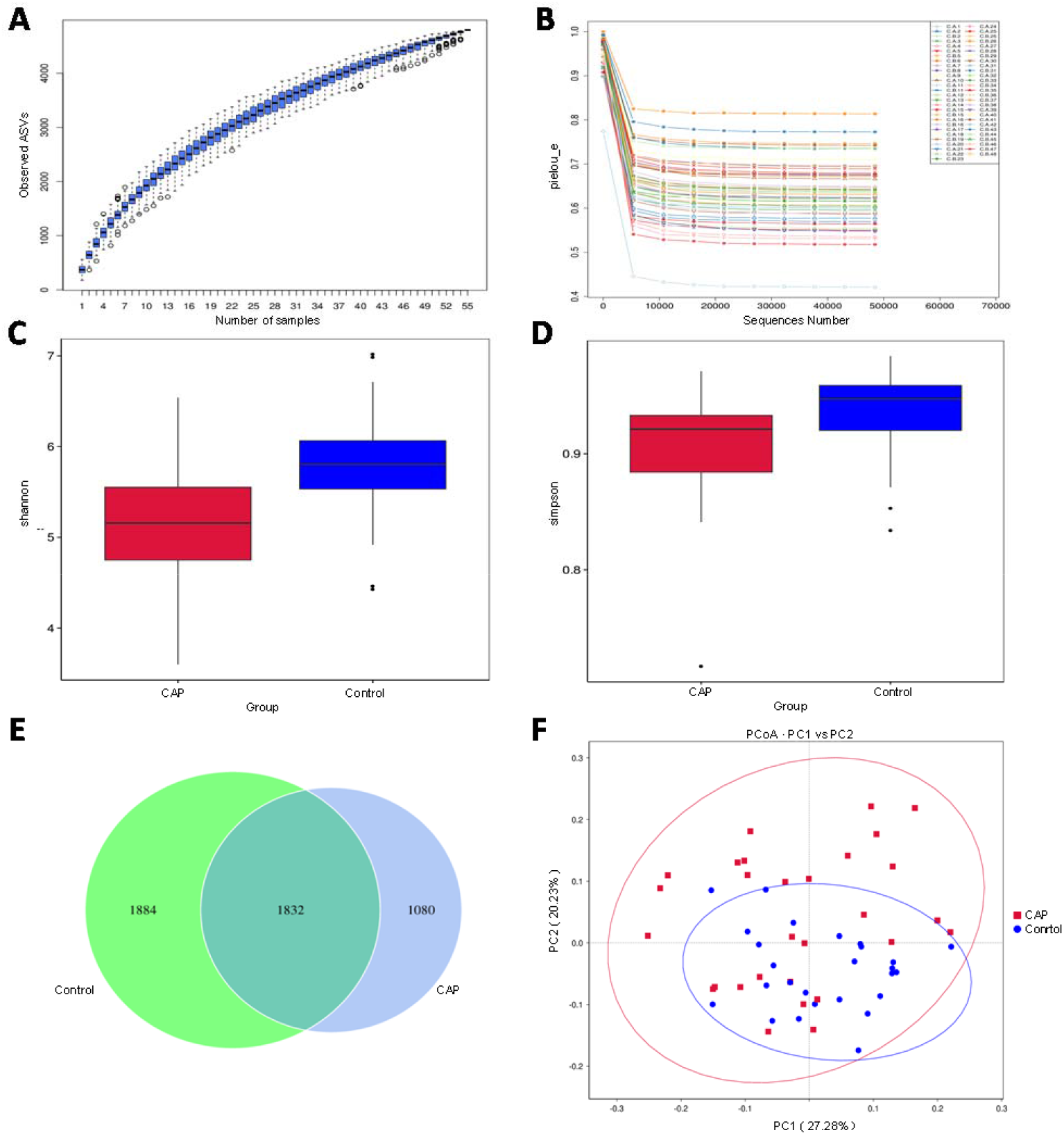
Bacterial diversity of the oral microbiota. (A) Species accumulation curve between number of samples. (B) The pielou’s evenness reflected the evenness of the distribution of species in the microbial community. Oral microbial diversity was estimated by the Shannon index (C) and Simpson index (D). (E) A Venn diagram displayed the overlaps between groups. (F) Beta diversity was calculated using weighted UniFrac by PCoA.

### 3.3 Oral microbial composition is different between people with CAP and healthy controls

The average composition of bacterial communities at the phylum and genus levels was shown in Figures 2A&2B, respectively *Proteobacteria, Firmicutes, Bacteroidota* were the three dominant bacterial phyla in the two groups (<u>Table S3</u>). At the phylum level, *Proteobacteria* was significantly lower in the CAP group than in the control group (Figure 2A). At the genus level, the dominant bacteria in the two groups were *Neisseria* and *Streptococcus*. Compared with the control group, *Neisseria* was significantly decreased, and *Prevotella* was significantly increased (Figure 2B, <u>Table S4</u>). The top 5 bacterial genera contributing to the difference between the two groups were *Neisseria, Streptococcus, Haemophilus, Prevotella, Fusobacterium* (Figure 2C-D, <u>Table S5</u>). At the genera level, *Prevotella, Veillonella, Campylobacter* were significantly enriched in the CAP group compared with the control group. And *Neisseria, Fusobacterium* were significantly reduced in the CAP group than in the control group (P < 0.05)(Figure 2E, <u>Table S6</u>). At the species level, *Prevotella_melaninogenica, Campylobacter_concisus* were significantly enriched in the CAP group compared with the control group. And *Fusobacterium_periodonticum, Prevotella_nanceiensis, Haemophilus_parainfluenzae* were significantly lower in the CAP group than in the control group (P < 0.05)((Figure 2F, <u>Table S7</u>).

**Figure 2.**
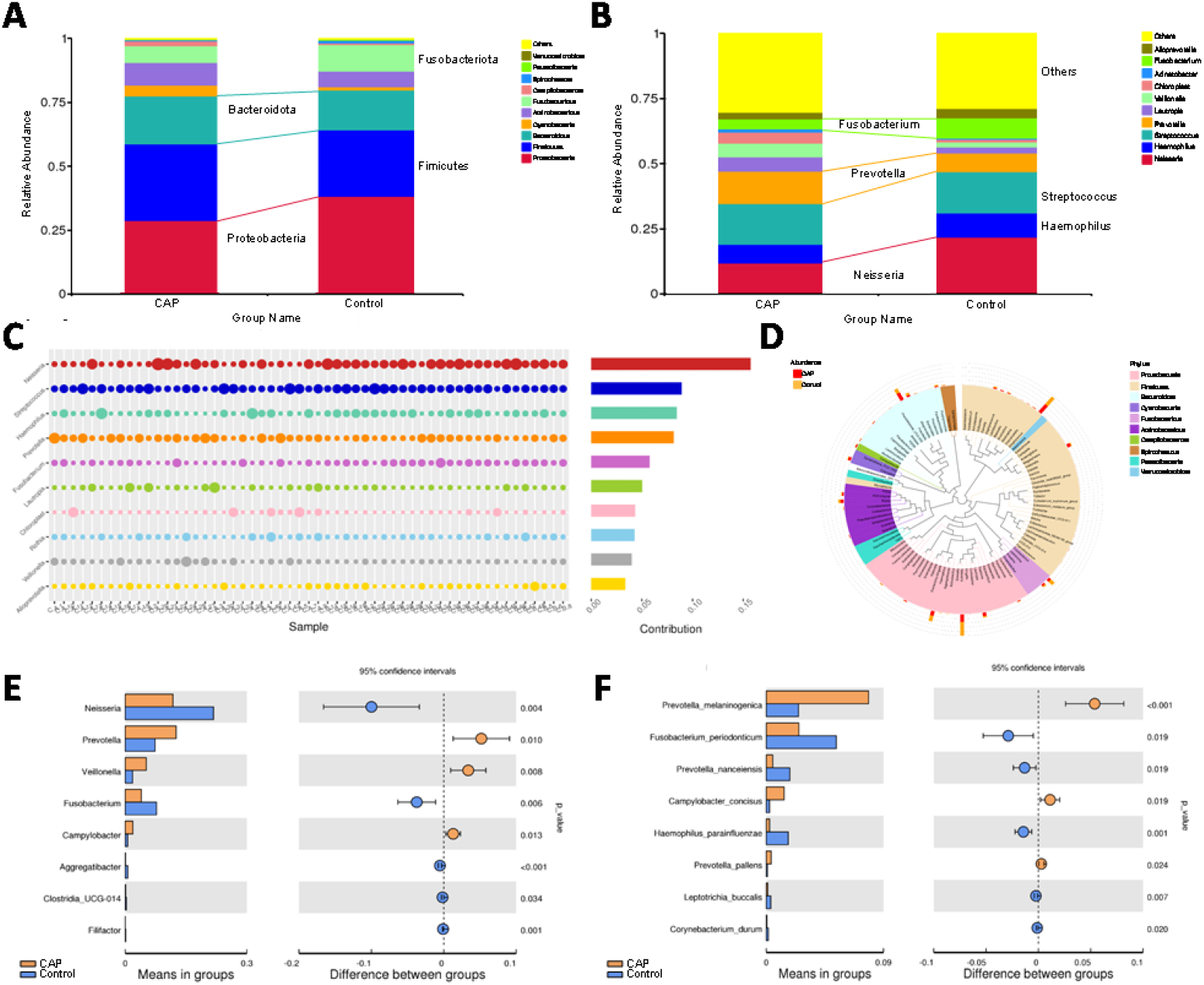
Oral microbiota composition. Average composition of bacterial community at the phylum (A) and family (B) levels. Abundance of the different bacterial genera (C) and species (D) between the CAP group and control group. The increased or decreased microbial community at the genus level (E) and species level (F) in patients with CAP versus healthy controls.

### 3.4 Oral bacteria are associated with infectious disease in people with CAP

To study the functional and metabolic changes in oral microbial communities, all ASVs were aligned into the Phylogenetic Investigation of Communities by Reconstruction of Unobserved States (PICRUSt) built-in reference database (15, 20). A principal component analysis was then performed using the total KEGG pathways data generated from all samples (Figure 3A). PICRUSt analysis identified 11 KEGG pathways with significant differential abundance between the CAP and control group (Figure 3B). As shown in Figure 3B, the pathway involving infectious disease was overrepresented in the CAP groups compared with the control group. Moreover, the pathways involving cell motility and energy metabolism were overrepresented in the CAP groups compared with the control group (figure 3C). The association between differential bacteria and metabolic pathways was investigated using correlation heatmaps. As the results showed, genera *Veillonella, Prevotella* and *Campylobacter* were positively correlated with infectious disease, while genera *Neisseria* and *Fusobacterium* were negatively correlated with the infectious diseases (Figure 3C-D).

**Figure 3.**
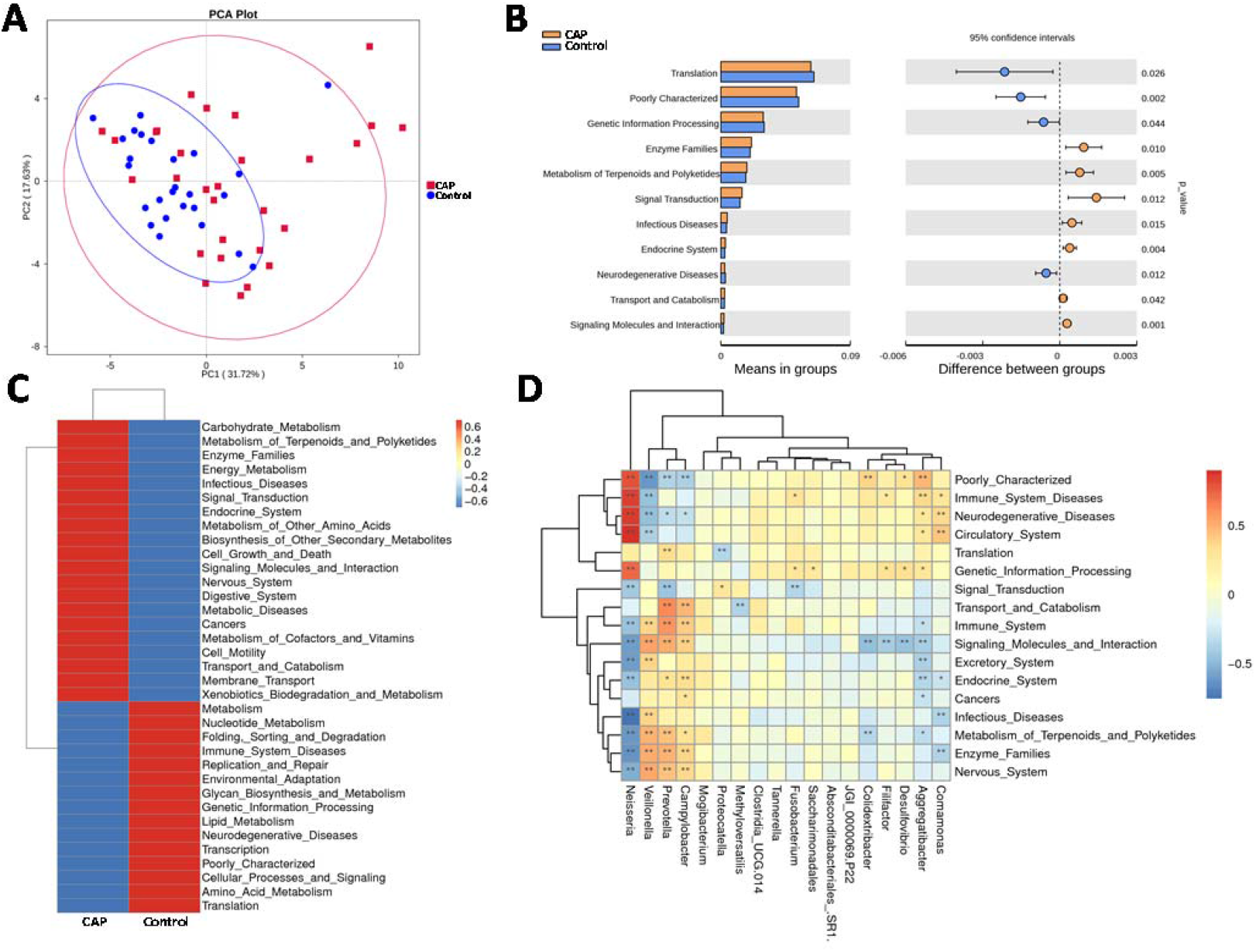
Oral microbial functional dysbiosis in patients with CAP and control group. Differential KEGG pathways were analyzed using PICRUSt, and PCA (A) analysis was conducted for the two groups. PICRUSt analysis identified 11 KEGG pathways (B) with significant differential abundance between the CAP and control group. The heatmaps (C—D) show partial spearman correlation coefficients of significant bacterial genera and differential KEGG pathways between CAP and control group.

## 4. Discussion

Management of CAP focuses particularly on early administration of the appropriate antimicrobial agent (22). Current diagnostic methods may be limited by sampling difficulties, delayed results, and difficulties in interpretation (5). Our research used the oral flora as the diagnostic factor of CAP, which effectively solves the problem of sampling difficulties. Our data discovered the significantly higher abundance of *Prevotella, Veillonella* and *Campylobacter*, and lower abundance of *Neisseria* and *Fusobacterium* than healthy group, which can be used as biomarkers in diagnosis of CAP.

The baselines of our study population were consistent. Specifically, the basic information and oral health status were not statistically different. We recruited the accompanying family members of the patients as a control group, and they could maintain good consistency in terms of living habits, eating habits, environment, socioeconomic conditions, etc. This can greatly reduce the interference of confounding factors on the oral flora and ensure the accuracy of the results.

The oral bacteria composition of patients with COPD also found the loss of microbiota diversity (23). We also found that CAP group had a lower diversity than healthy controls. Due to risk factors such as increased inhalation and mechanical ventilation under abnormal conditions, oral microorganisms may transfer to the respiratory tract and aggravate respiratory disease (24, 25).

*Prevotella* is consistently among the core bacterial communities of the respiratory tract (26, 27), which is known to be associated with pneumonia (14, 28, 29). Research found that the periodontopathic bacterial genera *Prevotella* were increased in the BALF of patients with CAP (30). We discovered an enrichment of *Prevotella*, especially *Prevotella_melaninogenica* in oral cavity of patients with CAP. It suggests that the mouth may be a reservoir of bacteria for the lungs. In a mouse lung co-infection model, the increased abundance of *Prevotella melaninogenica* was found, which probably was a potentially beneficial role to reduce infection (31). The above suggests that there was consistency in the changes of oral and pulmonary bacteria (*Prevotella* and *Prevotella_melaninogenica*) under lung inflammation.

We also observed that the abundance of *Veillonella* and *Campylobacter* were significantly higher in the CAP group. Few studies have focused on their relationship with CAP. But in a lung cancer model, lower airway dysbiosis with *Veillonella* led to decreased survival time and increased tumor burden (32). Skallevold et al. found that *-Veillonella* and *Streptococcus* in saliva could be valuable markers with diagnostic and prognostic potentials in lung cancer (33). On the other hand, in COPD, asthma, and cystic fibrosis (CF) lung diseases, oral microbiota (e.g. *Prevotella* species, *Veillonella* species) were present in the airways and associated with increased host inflammatory response (26-28). Additionally, Veillonella was identified as the most prominent biomarker for COVID-19 group sequencing analyses of oropharynx swab specimens (29, 34). Therefore, the potential of *Veillonella* and *Campylobacter* as a diagnostic biomarker for lung disease is proposed.

In control group, *Neisseria* and *Streptococcus* were the core genus of oral flora. The abundance of *Neisseria* and *Fusobacterium* was distinctly lower in CAP group than control group. Similarly, the levels of *Neisseria* in the pharynx were significantly lower in COPD patients compared to healthy people (35). Lower lung function, COPD diagnosis, and greater symptoms were recognized an association positively with *Neisseria* in lung microbiota (36). LaMotte et al. reported that oral *Neisseria spp*. could be causative bacteria in ventilator-associated pneumonia (VAP) patients (37). *Neisseria* is possibly an infection-promoting factor for lung disease. There are very few studies on the relationship between *Neisseria* and CAP, which is in urgent need of illustration.

*Fusobacterium*, recognized as a periodontal pathogen, is considered as a biomarker of lung function deterioration of COPD patients coinfected with *Pseudomonas aeruginosa* (38). Similar to oral flora changes in the present study, the *Fusobacterium periodonticum* resulted as the most significantly reduced species in COVID-19 patients (39). Consequently, *Fusobacterium*, especially *Fusobacterium periodonticum* was a potential specific detection indicator.

The functional analysis showed that the genera *Veillonella, Prevotella* and *Campylobacter* were positively correlated with infectious disease, while genera *Neisseria* and *Fusobacterium* were the negative correlations. The result further illustrated that these microbiotas can potentially serve CAP as biomarkers.

Anyway, our results provide baseline associations between oral microbiota and CAP, which will be important for further evaluation of treatment and prognosis.

## 5. Limitations

There are some limitations about this study. The sample size can be expanded, and we just conducted a single-center study instead of a multi-center research. Further exploration about the verification of biomarkers and potential mechanism involved in CAP are required in the future.

## 6. Conclusions

Oral microbial dysbiosis was found in patients with CAP. Dysbiosis of the oral microbiota (e.g. *Prevotella* and *Neisseria*) and neglected oral bacterial genera (e.g. *Veillonella, Campylobacter* and *Fusobacterium*) are important biomarkers for CAP.

## Supplementary Material

Supplementary tables S1-S7

## Abbreviations

CAP: community-acquired pneumonia
ASVs: amplicon sequence variants
PCoA: principal coordinate analysis
PICRUSt: phylogenetic investigation of communities by reconstruction of unobserved states
BALF: bronchoalveolar lavage fluid
VAP: ventilator-associated pneumonia
COPD: Chronic obstructive pulmonary disease COVID-19 Coronavirus disease 2019
CF: cystic fibrosis

## Author contribution

Study concept and design: TZ, NS; Specimen provider: NS, XZ, YH; Acquisition of clinical data: NS, XZ, YH; Data analysis and interpretation (including statistical analysis): NS, XZ; Drafting of the manuscript: NS. All authors approved the current version of this manuscript.

## Funding

Supported by the Fundamental Research Funds for the Central Universities (11621038) and the Science and Technology Program of Guangzhou (202201010401)

## Declaration of competing interest

The authors declare that they have no known competing financial interests or personal relationships that could have appeared to influence the work reported in this paper.

## Ethical approval

This study was approved by the Institutional Review Board of IRB of Jinan University (JNUKY-2022-002). The study was performed in accordance with the Helsinki Declaration and Rules of Good Clinical Practice. All participants signed written informed consents after the study protocol was fully explained.

## Acknowledgments

We thank Dr Xingdong Cai (the First Affiliated Hospital, Jinan University) for help in samples collection and thank Novogene Co. Ltd. for 16SrRNA sequencing. We also thank the generous volunteer subjects who enrolled in the study.

